# Protocols for circulating neutrophil depletion in neonatal C57Bl/6 mice

**DOI:** 10.1101/2024.05.06.592835

**Authors:** Devashis Mukherjee, Sriram Satyavolu, Sarah Cioffi, Asha Thomas, Yuexin Li, Lalitha Nayak

## Abstract

Murine neonatal neutrophil depletion strategies have problems achieving deep neutrophil clearance and accurate residual neutrophil fraction detection. An isotype switch method can achieve profound neutrophil clearance using a combination of anti-Ly6G and anti-rat κ Ig light chain antibodies in adult C57Bl/6 mice, proven by extra- and intracellular Ly6G detection by flow cytometry. We adapted this technique to neonatal mice, testing four neutrophil depletion strategies in the peripheral circulation, bone marrow, and spleen. Four protocols were tested: P3 Ly6G and P1-3 Ly6G (anti-Ly6G on postnatal days (P) 3 and 1-3 respectively), and P3 Dual and P1-3 Dual (anti-Ly6G and anti-rat κ Ig light chain on P3 and P1-3 respectively). Intracellular and extracellular Ly6G presence was detected using flow cytometry. Isotype control antibodies were used as controls. P1-3 Dual protocol achieved significantly better neutrophil depletion than the P1-3 Ly6G or P3 Ly6G protocols (97% vs. 74% and 97% vs. 50%, respectively) in the peripheral circulation. The P3 Dual protocol alone was enough to achieve significantly better neutrophil clearance (93%) than any of the Ly6G alone protocols. The Ly6G alone protocols led to near-total elimination of extracellular Ly6G. However, there was a significant presence of intracellular Ly6G in the CD45+ cell population, evading detection by extracellular Ly6G antibody-based detection methods. P3 protocols perform better than P1-3 protocols for bone marrow and splenic neutrophil clearance. Thus, the P3 Dual protocol might be the most effective and ethical protocol to induce profound neutrophil depletion in neonatal mice, an alternative to daily anti-Ly6G injections.

**Summary Sentence:** Dual antibody-based neutrophil depletion effectively induces circulating neutrophil clearance in neonatal mice.

## INTRODUCTION

Neonates, especially those born preterm, have a significantly higher risk of mortality and morbidity from sepsis than any other age group [1, 2]. This has been attributed to an early exit from a sterile intra-uterine environment to a microbe-laden external environment, a naïve adaptive immune system, and a developmentally immature innate immune system [3]. Neutrophils form the largest fraction of the innate immune system, and neonatal neutrophils are structurally and functionally different from adult neutrophils [4]. Elucidating neutrophil-dependent mechanisms contributing to increased sepsis-related mortality in neonates is critical in developing novel therapeutic strategies. Depletion of neutrophils is often used in murine models of sepsis to characterize the role and contribution of the circulating neutrophil compartment. Extrapolating adult neutrophil depletion methods to neonatal mice is not recommended, as neonatal neutrophil stores, production, and turnover are distinct from those at other ages. Neonatal neutrophils have been described as resistant to apoptosis, and neonates have a diminished bone marrow reservoir of precursors with a limited ability to mobilize them in response to infection.[4, 5] Unlike a steady-state circulating neutrophil population in adults, neonates have a fluctuating neutrophil count, and these neutrophils have a prolonged survival time in circulation.[6, 7] Thus, it is critical to test neutrophil depletion protocols in neonatal models to identify the least invasive and most efficacious strategy that minimizes the number of painful injections at this early age.

Neutrophil depletion strategies include pharmacologic agents such as cyclophosphamide and vinblastine and antineutrophil antibodies [8, 9]. Pharmacologic neutrophil depletion has fallen out of favor due to effects on other cell lines and other cellular processes such as microtubule assembly [10]. Murine models that use antibody-mediated neutrophil depletion use either anti-Ly6G or anti-Gr1 antibodies [9]. Anti-Gr1, RB6-8C5 clone is a rat IgG2b, which leads to neutrophil depletion by complement-mediated cell cytotoxicity, in addition to the classical pathway, and is thus faster and more pronounced [11]. However, the anti-Gr1 antibody also recognizes Ly6C receptors, expressed by monocytes, macrophages, and a subpopulation of CD8+ T cells [9, 12]. This leads to the non-specific depletion of several different cell types [13]. Anti-Ly6G antibody is a rat IgG2a and binds specifically to the Ly6G receptors on neutrophils and leads to neutrophil depletion by Fc-opsonization and macrophage phagocytosis, which is a slower method than the complement-mediated cytotoxicity seen in anti-Gr1 depletion [11]. Boivin et al. have developed a new model for sustained and more durable depletion of neutrophils in adult mice where they injected anti-Ly6G daily and anti-rat IgG2a antibody every other day [11]. The authors mention that this “Combo” treatment creates an isotype switch in which the anti-Ly6G depleting antibody (rat IgG2a isotype) is bound by the anti-rat IgG2a (mouse IgG2a isotype). Mouse IgG2a is orthologous to rat IgG2b, thus creating an isotype switch and leading to faster and more profound neutropenia by complement-mediated processes. This group circumvented the issue of antigen masking during the detection of the residual circulating neutrophils to measure the efficacy of neutrophil depletion. Although extracellular Ly6G staining would detect circulating neutrophils using flow cytometry under standard conditions, those circulating neutrophils to which the anti-Ly6G depleting antibodies are bound are not detected by this technique due to antigen masking. This hurdle was overcome using a combination of extracellular Ly6G staining, intracellular Ly6G staining, and extracellular rat IgG2a staining [11]. Using this method, rat IgG2a staining can detect those residual circulating neutrophils still bound to the anti-Ly6G depleting antibody and evade detection by fluorophore-conjugated anti-Ly6G antibodies.

Neonatal neutrophil depletion studies commonly use anti-Ly6G depleting antibodies to study the role of the neutrophil [14-16]. These studies generally report >90% neutrophil depletion using fluorophore-conjugated extracellular Ly6G detection on flow cytometry, which does not detect the circulating neutrophils bound to the depleting antibody. Data on the accurate degree of neutrophil depletion using extra- and intracellular staining methods are absent. In addition, no studies have replicated the isotype switch in neonatal mice to demonstrate whether this leads to more profound neutrophil depletion or whether an isotype switch model is unnecessary in neonatal mice. Here, we describe a modified Boivin neutrophil depletion strategy for postnatal day 4 (P4) neonatal mice and evaluate the efficacy of four protocols in the peripheral circulation, bone marrow, and spleen. Further, we tested the combined staining method for evidence of neutrophil depletion in the conventional anti-Ly6G depletion and the Boivin “Combo” depletion strategy, adapted for neonatal mouse pups.

## MATERIALS AND METHODS

### Animals

All animal experiments were conducted following the approved protocols of the Case Western Reserve University (CWRU) Institutional Animal Care and Use Committee. Mice colonies were housed in a pathogen-free animal facility. C57Bl/6 mice procured from The Jackson Laboratory (Bar Harbor, Maine, USA) were used for all experiments. Neonatal mice were anesthetized using isoflurane following CWRU Animal Resource Center protocols for peripheral blood sampling from the external jugular vein. Pups were euthanized after extraction of blood using cervical dislocation as per CWRU and NIH guidelines for rodents < 10 days of age.

### In vivo antibodies

The anti-Ly6G antibody (clone 1A8, #BP0075-1), the anti-rat kappa immunoglobulin light chain antibody (clone MAR18.5, #BE0122), and the corresponding isotype controls were procured from BioXCell (Lebanon, NH, USA). Anti-Ly6G was used at a dose of 100ug/gm, and anti-rat k IgG light chain antibody was used at a dose of 50ug/gm. Intraperitoneal injections were administered to mice using 31-gauge syringes.

### Neutrophil depletion protocols (Fig. 1A)

One-day anti-Ly6G (P3 Ly6G) – Anti-Ly6G antibody injection on P3, followed by peripheral blood (PB) sampling on P4 after 24 hours (h).

**Figure 1.**
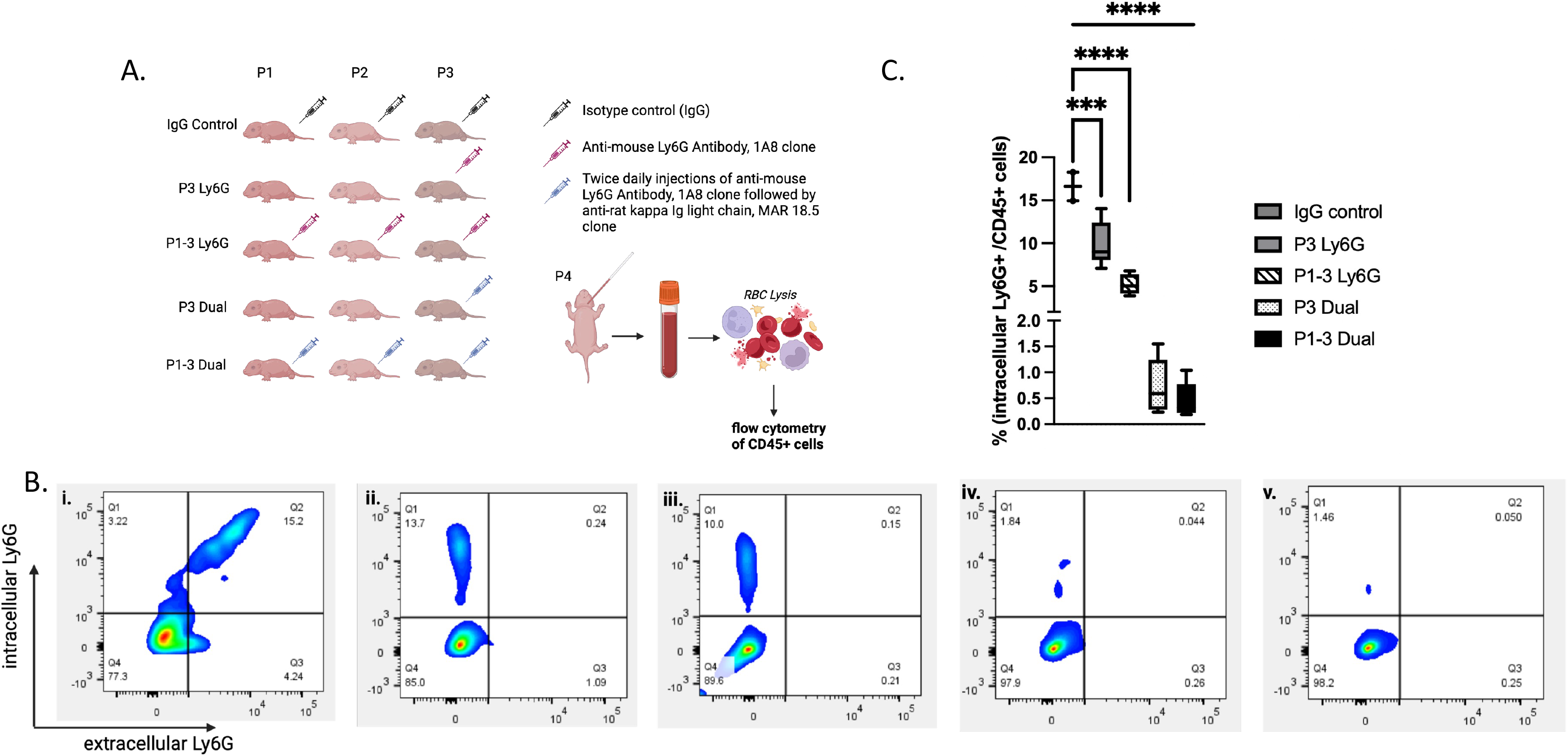
(A) P3 protocols included intraperitoneal injections administered on postnatal day three only, and P1-3 protocols included intraperitoneal injections administered daily on postnatal days 1-3. All protocols were followed by peripheral blood sampling on P4 24 hours after the last injection. Red blood cell lysis was performed using hypotonic saline, reconstituted to normal saline, and cells were resuspended in flow cytometry buffer. Extracellular staining was performed by APC-CD45, FITC-Ly6G (1:1000), cells were fixed and permeabilized, and intracellular staining was performed by PE-Ly6G (1:1000). (B) Flow cytometry was performed on BD LSF Fortessa cytometer for all protocols (i) IgG control (n=2), (ii) P3 Ly6G (n=5), (iii) P1-3 Ly6G (n=4), (iv) P3 Dual (n=6), and (v) P1-3 Dual (n=5). We included quarters 1 – 3 for a total neutrophil count (CD45+ cells staining either intracellularly or extracellularly for Ly6G). As compared to the mean neutrophil count in isotype control injected mice, P3 Ly6G mice exhibited ∼50% reduction, P1-3 Ly6G mice showed 74% reduction, P3 Dual mice showed 93% reduction, and P1-3 Dual mice exhibited 97% reduction in the circulating neutrophil counts. (C) Percentage of intracellular Ly6G positive cells out of all CD45+ cells. As compared to those in isotype control injected mice (16.6 ± 2.34), this percentage decreased only by 6.63% in P3 Ly6G injected mice and by 11.41% in P1-P3 Ly6G injected mice (p<0.0001). However, this reduction was far more effective in both protocols involving anti-rat kappa Ig light chain antibodies. The percentage of CD45+ cells positive for intracellular Ly6G reduced by 15.86% in P3 Dual mice (p<0.0001) and by 16.14% in P1-3 Dual mice (p<0.0001), thus showing the effectiveness of the dual antibody treatment protocols in preventing antigen masking and Ly6G receptor internalization.

Three-day anti-Ly6G (P1-3 Ly6G): Daily anti-Ly6G antibody injections were administered from P1 to P3, followed by peripheral blood (PB) sampling on P4 24 h after the last injection. One-day dual-antibody treatment (P3 Dual) – Anti-Ly6G and anti-rat kappa immunoglobulin light chain antibody on P3, followed by PB sampling after 24 h.

Three-day dual-antibody treatment (P1-3 Dual) – Daily anti-Ly6G and anti-rat kappa immunoglobulin light chain antibody injections from P1 – P3, followed by PB sampling on P4, 24 hours after the last injection. Anti-Ly6G and anti-rat k IgG light chain antibody injections were spaced out by a four-hour interval.

Peripheral blood procurement – The left facial vein was accessed in anesthetized P4 pups using a previously described method.[17] Briefly, under microscopic visualization, the left earbud was located, and the area around it was sterilized with 70% ethanol. The facial vein runs anterior to the earbud and is visible through the skin in P4 pups. It increases in prominence after wiping the area with ethanol. After careful dissection of the overlying skin, the facial vein was isolated and accessed to obtain up to 20uL of peripheral venous blood.

Bone marrow and spleen isolation – Bilateral femurs and tibias of anesthetized P4 pups were isolated under sterile conditions, crushed under a laminar flow hood, and passed through a 70um cell strainer to obtain a live cell suspension of bone marrow cells. A similar technique was employed for splenic cells to obtain a live cell suspension.

Red blood cell lysis – All cell suspensions were resuspended in 10mL of hypotonic 0.2 % w/v (0.034 M) NaCl for 30 seconds to lyse red blood cells. This was then mixed with an equal volume of 1.6 % w/v (0.274 M) NaCl to reconstitute the cell suspension to a final concentration of 0.9% w/v (0.154 M) isotonic saline. This was centrifuged at 350 x g at 4 °C for 10 minutes, and the cell pellet was resuspended in cell staining buffer as described below.

<u>Extra- and intracellular</u> staining protocol – All steps were protected from light. Isolated cells were resuspended in fluorescent activated cell sorting (FACS) buffer (phosphate-buffered saline (PBS) + 1% fetal bovine serum + 2mM EDTA), incubated with cell viability stain (Live/Dead Fixable Yellow Dead Cell Stain kit, Life Technologies, USA) for 20 minutes, washed in FACS buffer for 5 minutes at 350 x g and resuspended in 100uL of FACS buffer. The cells were then incubated with an FcR blocker (anti-mouse CD16/32, BioLegend #101319) for 10 minutes on ice to block the non-specific binding of antibodies to Fc receptors. For extracellular staining, cells were incubated on ice with fluorescent-conjugated antibodies – anti-mouse-CD45-APC (BioLegend) and anti-mouse-Ly6G-FITC (eBioscience) at a concentration of 1:100 for 30 minutes, washed with 1mL of FACS buffer, centrifuged at 350 x g for 10 minutes, and fixed immediately for intracellular staining. For fixation, cells were incubated in 250uL of fixation buffer (BioLegend FluoroFix buffer) at room temperature for 1 hour. Fixed cells were permeabilized using BioLegend Permeabilization buffer by incubating in 100uL of the buffer on ice for 20 minutes, followed by two 5-minute washes using permeabilization buffer at 350 x g.

Permeabilized cells were re-fixed using the fixation buffer for 5 minutes at room temperature, followed by incubation with anti-mouse-Ly6G-PE antibody at 1:100 in permeabilization buffer for 30 minutes at room temperature for intracellular Ly6G staining. This was followed by two 5-minute washes using the permeabilization buffer at 350 x g. Cells were then resuspended in 100uL of FACS buffer and processed for flow cytometry within 24 hours.

Flow cytometry - All flow cytometry experiments were conducted using BD LSR Fortessa cytometer at the CWRU Cytometry Core. Gating strategies included multi-color compensation using unstained controls and antibody capture compensation beads (BD CompBeads Compensation Particles) using the same antibodies described above. The gating tree included forward and side scatter (FSC/SSC), followed by live cell gating (cells negative for live/dead stain), followed by SSC/APC-CD45 (the entire leukocyte population), and followed by PE/FITC for intra- and extracellular Ly6G expression respectively. A total of 50,000 cells were counted for each sample. Each experiment included unstained controls to ensure no background fluorescence was detected. All axes were in logarithmic scales. Flow cytometry results were analyzed, and representative images were built using FlowJo version 10.8 (BD Biosciences, Oregon, USA).

### Statistical analysis

GraphPad Prism was used for all statistical analyses, with significance considered when p <0.05. Standard one-way ANOVA was used to test differences in means of all protocols. A standard unpaired t-test with Welch’s correction was used to test differences in means between each protocol and the control IgG protocol.

## RESULTS

### Administering Anti-Ly6G antibody alone does not induce profound neutrophil depletion in P4 pups

We found no difference between previously reported circulating neutrophil counts in neonatal C57Bl/6 mice [18], and the circulating fraction of CD45+ cells in P4 PB, which were positive for either intracellular or extracellular Ly6G or both after isotype control injections in our studies (Fig. 1B (i) and Table 1). Both P3 Ly6G and P1-3 Ly6G protocols effectively removed any extracellular presence of Ly6G on circulating CD45+ cells, to 1.3% and 0.3%, respectively, from a baseline level of 19.6% (Fig. 1B). Using only extracellular Ly6G-based detection techniques to detect residual neutrophils would thus show >98% neutrophil depletion using either protocol. However, using intracellular Ly6G staining, we demonstrate that the P3 Ly6G protocol increased antigen masking by increasing the fraction of cells with only internal Ly6G expression and no detectable extracellular Ly6G expression from 2.1% to 9.67% (p = 0.002) (Fig. 1 B and C). P1-3 Ly6G leads to fewer neutrophils with only intracellular Ly6G expression (5.08%), but it still did not achieve >98% neutropenia based on intracellular + extracellular Ly6G detection (Fig. 1 B and C). In our studies, both P3 Ly6G and P1-3 Ly6G protocols achieved ∼50% and ∼74% neutrophil depletion, contrary to what has been reported previously using only extracellular Ly6G detection. We use fluorophore-conjugated anti-rat IgG2a antibodies to detect the circulating neutrophils to which anti-Ly6G antibodies were already bound, and this ranged between 10-12% of all CD45+ cells in both P3 Ly6G and P1-P3 Ly6G protocols, despite both protocols showing <2% extracellular Ly6G+ cells, further proving that the depleting antibody itself impairs detection of the Ly6G receptor, similar to what Boivin et al. reported.

**Table 1:** Peripheral blood circulating cells expressed as a percentage of live CD45+ cells positive for either intracellular or extracellular Ly6G expression and those cells that express Ly6G both intra- and extracellularly. The rightmost column gives the neutrophil counts for each protocol based on whether the cell expressed Ly6G (intra- or extracellular).

|  | Only intracellular Ly6G expression (% of CD45+ cells) | Only extracellular Ly6G expression (% of CD45+ cells) | Both intra- and extracellular Ly6G expression (% of CD45+ cells) | Total neutrophil count (% of CD45+ cells) | Neutrophil depletion percentage based on isotype control arm |
| --- | --- | --- | --- | --- | --- |
| Isotype control (n=2) | 2.10 ± 0.64 | 3.40 ± 0.60 | 14.50 ± 1.70 | 20.00 ± 2.93 | 0% |
| P3 Ly6G (n=5) | 9.67 ± 2.62 | 0.36 ± 0.48 | 0.17 ± 0.25 | 10.32 ± 3.02 | 50% |
| P1-3 Ly6G (n=4) | 5.08 ± 1.09 | 0.09 ± 0.06 | 0.11 ± 0.13 | 5.28 ± 1.15 | 73% |
| P3 Dual (n=6) | 0.50 ± 0.46 | 0.75 ± 0.49 | 0.23 ± 0.27 | 1.49 ± 0.63 | 93% |
| P1-3 Dual (n=5) | 0.42 ± 0.35 | 0.173 ± 0.09 | 0.04 ± 0.03 | 0.63 ± 0.30 | 97% |

### P1-3 Dual protocol induces more significant neutrophil depletion than P1-3 Ly6G protocol

Adapting the established “Combo” depletion strategy for neonatal mice, we show that dual antibody treatment with anti-mouse-Ly6G followed by anti-rat IgG2a leads to more profound neutrophil depletion (p<0.0001) (Fig 1B and Table 1). Unlike Boivin et al.’s methodology of every other day anti-rat IgG2a antibody, we injected the mouse pups with both antibodies daily, separated by 4-6h. Our primary rationale was that, unlike adult experiments where cell depletion can be carried out for days to weeks before endpoint experiments, we only had three days to produce neutrophil depletion in neonatal pups and wanted to maximize the effect of the depleting antibodies. P1-3 Dual produced ∼97% neutrophil depletion in P4 mouse pups based on intracellular and extracellular Ly6G detection methods (Table 1). The fraction of CD45+ cells expressing only intracellular Ly6G, typically evading detection, was 0.4% (Fig. 1B (v) and Fig. 1C).

### P3 Dual protocol is a better alternative than P1-3 Ly6G and P3 Ly6G protocols for circulating neutrophil depletion

Administering two i.p. injections daily to neonatal mice can be challenging. We modified our approach to only P3 injections 24 hours before PB sampling. The P3 Dual protocol induced more profound neutrophil depletion than the P1-3 Ly6G protocol (∼93% vs. ∼74%, p=0.0074), the more commonly reported protocol for neonatal neutrophil depletion. This significant reduction in circulating neutrophil count was primarily attributable to a decrease in the neutrophils with intracellular Ly6G expression, similar to the results in the P1-3 Dual protocol (Fig 1B (iv) and Fig. 1C). P3 Dual is more effective than P1-3 Ly6G, as expected, in depleting cells expressing Ly6G intra- or extracellularly or both (Table 1).

### P3 regimens are more effective at bone marrow and splenic neutrophil clearance than P1-3 regimens

Although circulating neutrophils respond to sites of inflammation or infection, it is critical to evaluate residual neutrophil fractions in other primary neutrophil reservoir sites. We evaluated the four regimens in bone marrow and spleen with intra- and extracellular Ly6G detection. Similar to peripheral circulation, all four regimens reduced extracellular Ly6G expression from a baseline of 60% in the bone marrow and 12.4% in the spleen in control IgG-treated pups to less than 5% in both the bone marrow and spleen in every regimen (Fig. 2A-F, 3A-F). However, when accounting for the total residual neutrophil population (any Ly6G expression), P3 Dual had the lowest mean counts compared to the other three regimens for bone marrow and splenic compartments (Fig. 2H, 3H). Interestingly, the P1-3 Ly6G regimen had a higher mean residual neutrophil count than the P3 Ly6G regimen in the bone marrow (45.1 vs. 37.9 percent of all CD45+ cells, p=0.0055) and the spleen (8.2 vs. 5.8 percent of all CD45+ cells, p=0.0469). Similarly, the P1-3 Dual regimen also had a higher mean residual neutrophil count than the P3 Dual regimen in the bone marrow (36.5 vs. 21.7 percent of all CD45+ cells, not significant) and in the spleen (9.0 vs. 3.9 percent of all CD45+ cells, p0.0427). The P1-3 Ly6G and the P1-3 Dual regimens could not significantly decrease residual neutrophil counts in the spleen (Fig. 3H). Overall, P3 Dual was able to deplete neutrophils the most consistently out of all the regimens in the bone marrow and the spleen.

**Figure 2.**
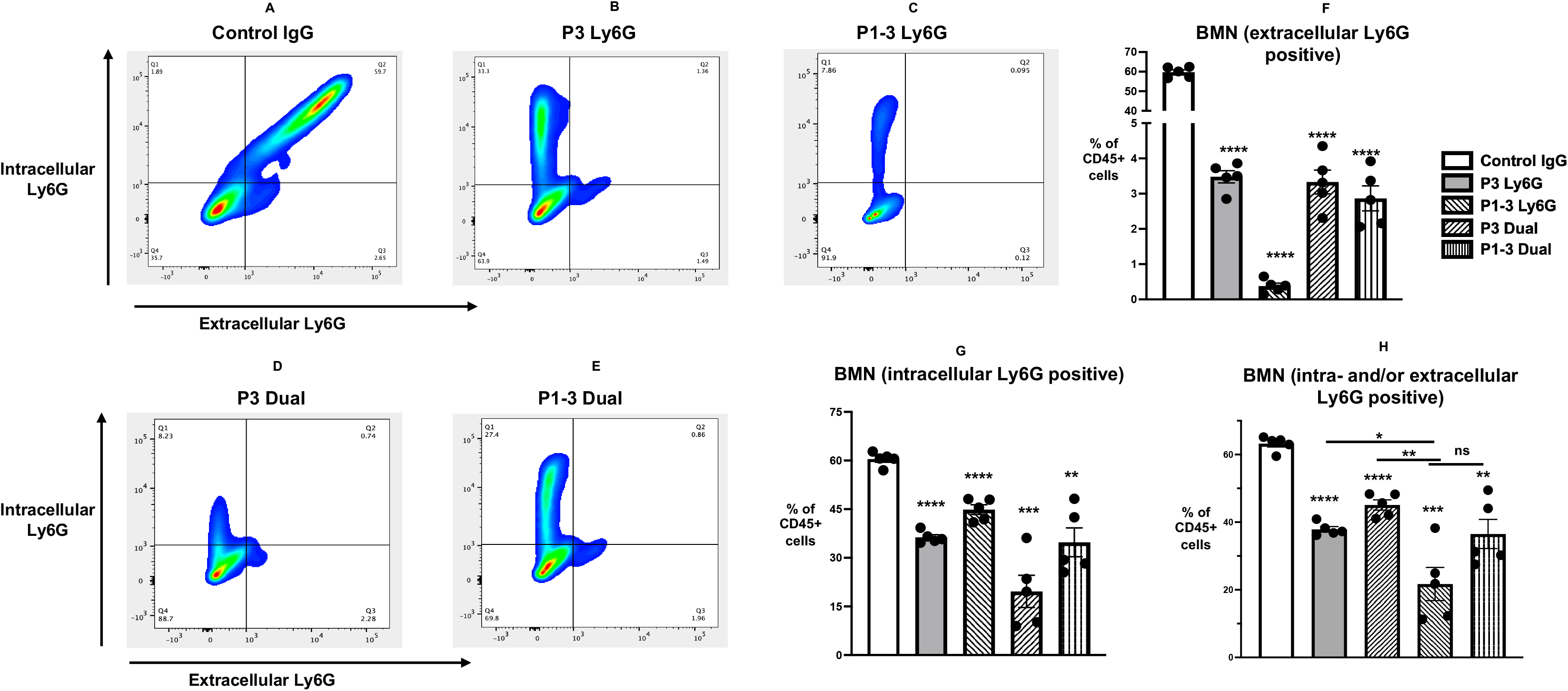
(A) – (E) show representative cytometry images from live, CD45+ bone marrow cells for intracellular (left y-axis, PE) and extracellular (x-axis, FITC) Ly6G expression for control IgG, P3 Ly6G, P1-3 Ly6G, P3 Dual, and P1-3 Dual neutrophil depletion protocols respectively. Fifty thousand events were counted for each experiment, and five biological replicates were used for each protocol. All protocols are effective at significantly eliminating extracellular Ly6G+ positive CD45+ cells from a baseline >60% (control IgG) to <10% (Fig. 2F). Although this decrease was most prominent in P3 Ly6G protocol, the differences in means of residual Ly6G expression between the four protocols is minimal at 2%. When counting all CD45+ cells positive for any Ly6G expression, the maximum decrease in residual neutrophil count is in the P3 Dual protocol (Fig. 2H), and this is likely from a significant decrease in the residual intracellular Ly6G positive cells in the P3 Dual protocol (Fig. 2G). The differences between means of residual neutrophil counts between the P3 Dual and the P1-3 Dual protocols were not significant. An unpaired t-test with Welch’s correction was used to analyze the statistical difference between the means of control IgG protocol and each protocol. * : p<0.05, ** : p<0.01, *** : p<0.001, **** : p<0.0001, ns not significant.

**Figure 3.**
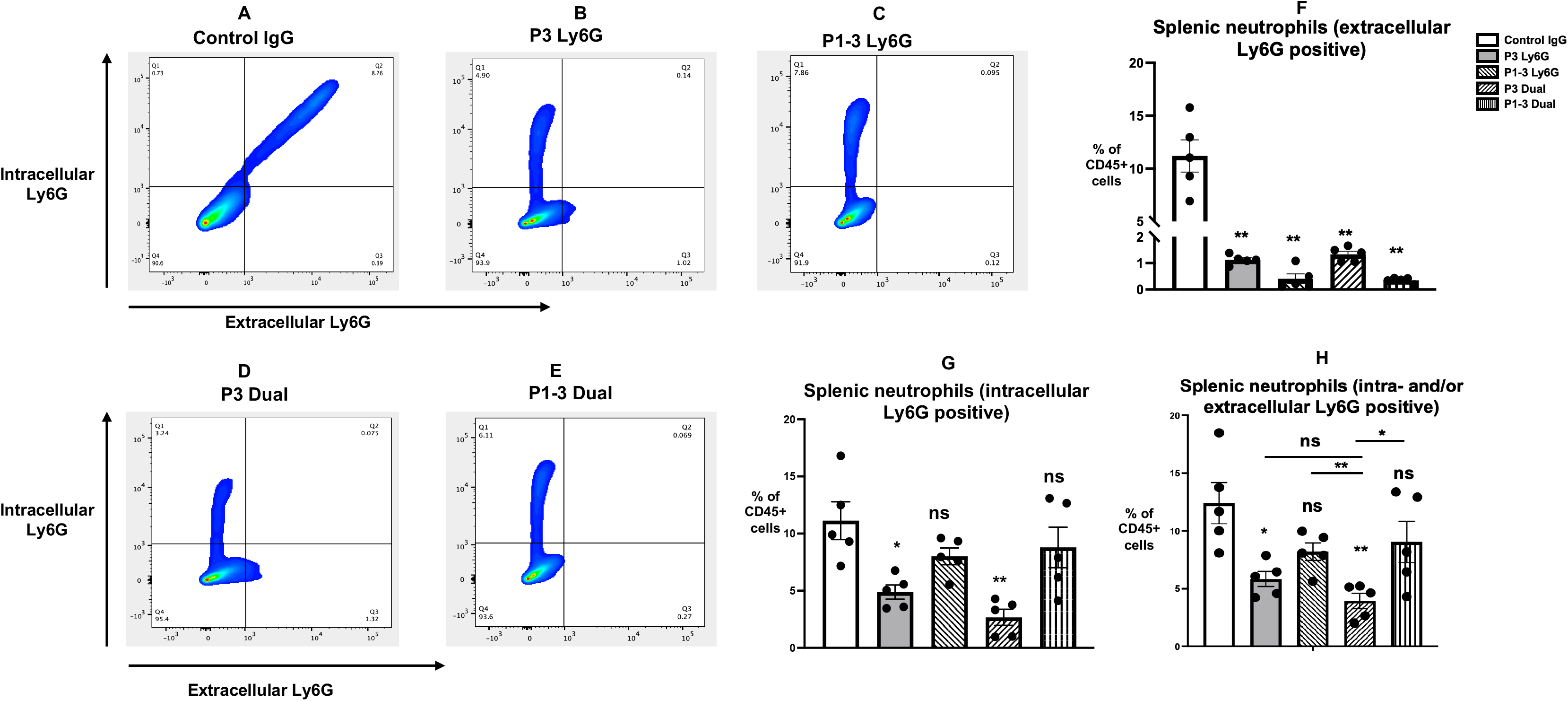
(A) – (E) show representative cytometry images from live, CD45+ splenic cells for intracellular (left y-axis, PE) and extracellular (x-axis, FITC) Ly6G expression for control IgG, P3 Ly6G, P1-3 Ly6G, P3 Dual, and P1-3 Dual neutrophil depletion protocols respectively. Fifty thousand events were counted for each experiment, and five biological replicates were used for each protocol. All protocols are effective at significantly eliminating extracellular Ly6G+ positive CD45+ cells from a baseline >10% (control IgG) to <50% (Fig. 3F). Both the P3 Ly6G and P3 Dual protocols were able to decrease the intracellular Ly6G positive fraction in the spleen significantly, but the P1-3 regimens were unable to demonstrate a statistically significant decrease (Fig. 3G). This effect is re-demonstrated when counting all residual neutrophils in the spleen (intra- and/or extracellular Ly6G positive cells out of all CD45+ live cells), where the P3 protocols can decrease the residual neutrophil count significantly, but the P1-3 protocols were unable to do so (Fig. 3H). The differences in means of residual neutrophil counts between the P3 Ly6G and the P3 Dual protocol were not significant. An unpaired t-test with Welch’s correction was used to analyze the statistical difference between the means of control IgG protocol and each protocol. * : p<0.05, ** : p<0.01, ns : not significant.

## DISCUSSION

These results show that a single dose each of the anti-Ly6G and anti-rat kappa immunoglobulin light chain antibodies administered 24 hours before is sufficient in inducing >90% circulating neutrophil depletion in neonatal pups, along with a significant reduction in neutrophil counts in the bone marrow and spleen. Our study is also the first to report on residual neutrophil counts in the BM and spleen after using the established anti-Ly6G regimens or the Dual antibody regimens. Our findings also show that unlike aged mice, where 20 days of dual antibody injections are necessary to induce profound circulating neutrophil depletion, neonatal neutrophil production and kinetics are different, and a single dose of both is sufficient. In addition, we uncovered an interesting finding that either P3 Ly6G or P3 Dual can similarly significantly reduce neutrophil counts in the spleen. However, the three-day regimens (P1-3 Ly6G or P1-3 Dual) could not significantly reduce the splenic neutrophil counts, although they achieved a similar reduction of extracellular Ly6G expression as seen with any of the regimens at any site. This is similar to previously published data where daily injections of anti-Ly6G or both anti-Ly6G and anti-rat antibodies could not achieve neutrophil depletion in the spleen.[11] Adapting an existing adult murine protocol described by Boivin et al. for neonatal studies adds significant knowledge to neonatal neutrophil biology. Neonatal neutrophil kinetics are unique and different from adults. Neonates have a decreased bone marrow storage pool of neutrophil precursors and mature neutrophils.[4, 19] Unlike adults, they cannot rapidly increase the turnover of neutrophil production, as the cell cycle from hematopoietic progenitor cell to a mature neutrophil is already maximally shortened in neonates. Thus, it is not possible to shorten this time further for the production and release of mature neutrophils.[20] After birth, the progenitor cell population is relatively small but grows rapidly in the next few days, and the ratio of mature neutrophils to progenitor cells falls quickly in the first week of life in rodent models.[21] Thus, it is critical not to extrapolate adult neutrophil depletion studies to neonatal models, especially as several days of injections are not feasible when the experimental endpoint is within the first week of life, and daily injections in neonatal pups significantly increase the ethical concerns associated with those studies.

Neutrophil depletion models are fraught with the problems of accurate and specific neutrophil population estimation after antibody treatment and the development of antibodies against the depleting antibody. The latter is not a problem with neonatal experiments, as mice take ∼seven days to develop anti-rat antibodies [11]. The production of the anti-Ly6G 1A8 clone antibody has improved the specificity of neutrophil depletion experiments, and most current studies use this antibody instead of the anti-Gr1 antibody, which also recognizes Ly6C expressed by other myeloid cells. Neonatal studies employing neutrophil depletion techniques use daily anti-Ly6G injections, and most of these studies have shown near-total circulating neutrophil depletion. These studies identify the residual neutrophil population by extracellular Ly6G staining, which runs the risk of incorrect estimation of neutrophils as they cannot detect the antigen-masked population. In our study, using the reported technique of dual Ly6G detection (intra-+ extracellular), we show that the previously published strategy of using daily anti-Ly6G antibody injections to induce neutrophil depletion in P4 neonatal mice leads to only ∼75% circulating neutrophil depletion, instead of the reported >90% depletion [14-16]. In addition, this cannot reduce splenic neutrophil counts; in the BM, neutrophil counts are reduced by 28%. Although anti-Ly6G antibodies can effectively remove extracellular Ly6G+ cells from circulation, neutrophils can internalize the Ly6G receptor as an adaptive mechanism against depleting antibodies, thus exhibiting decreased surface expression of the Ly6G receptor, preventing effective depletion by anti-Ly6G antibody treatment alone [22]. The increased time in depletion by only anti-Ly6G-based protocols may allow for this internalization to occur and lead to efflux of neutrophil precursors from the bone marrow, which might not have robust Ly6G expression extracellularly [23]. In addition, anti-Ly6G antibody leads to surface antigen masking and a false reassurance that neutrophil depletion has been achieved if only extracellular Ly6G expression is measured to detect residual neutrophils. Our results demonstrate this effect in all three sites, where the extracellular expression of Ly6G is almost non-existent compared to isotype control IgG, regardless of the choice of depleting antibodies or duration of treatment.

Boivin et al. have demonstrated that by creating an isotype switch from rat IgG2a (anti-Ly6G, 1A8), which is orthologous to mouse IgG1 to mouse IgG2a (anti-rat IgG2a, k light chain), faster and durable neutrophil depletion can be achieved [11, 24]. IgG2 subclasses activate both the classical and complement systems for neutrophil killing and thus are more effective and faster than IgG1-based depletion techniques. We show that a single dose of dual antibody treatment on P3 leads to more pronounced neutrophil depletion than three days of anti-Ly6G antibody treatment. This is particularly useful as injecting P1 mouse pups can be technically challenging and avoids daily injections in neonatal mice. In addition, at P4, 20% of the circulating CD45+ cells, 60% of the BM CD45+ cells, and ∼12% of the splenic CD45+ cells are neutrophils compared to 5-7%, ∼40%, and ∼1.2% respectively in 10-12-week-old mice. Despite having more neutrophil counts, using just P3 Dual, we were able to achieve significantly more neutrophil depletion in the peripheral blood compared to either P3 Ly6G or P1-3 Ly6G, which is in contrast to the adult data where one day of dual antibodies achieves the same peripheral blood depletion as one day of just anti-Ly6G alone or anti-Gr1 alone.[11] P3 Dual protocol achieved a similar decrease in the BM and spleen neutrophil populations as the adult one-day combination treatment, which was significant. However, interestingly, we were able to achieve significant splenic neutrophil depletion with either P3 Dual or P3 Ly6G, whereas in adult mice, only dual antibody treatment was able to accomplish this. Combining these results, we believe that the most efficacious dosing strategy for neonatal neutrophil depletion (including peripheral blood, BM, and spleen) is the P3 Dual regimen. The novelty of our technique lies in adapting this approach with modifications in the dosing schedule specifically for neonatal mice, which demonstrates highly effective neutrophil clearance compared to prior protocols.

As expected, the P1-3 Dual regimen achieved more neutrophil depletion in the periphery than the P3 Dual regimen (97% vs. 93%). However, the experimental relevance of this additional 4% depletion is not significant. The unexpected findings of this study were the less potent neutrophil depletion in both the BM and spleen achieved by either of the P1-3 regimens compared to their P3 counterparts (Figs. 2 and 3). As shown previously, administering neutrophil-depleting antibodies leads to the induction of granulopoiesis within one day.[11, 25] We hypothesize that the 3-day regimens lead to sufficient induction of granulopoiesis, which offsets the decrease in the neutrophil count achieved by the depleting antibodies, but this induction is not as prominent in the 1-day regimens. The circulating neutrophil counts are unaffected as there are significant circulating levels of depleting antibodies, which can neutralize the neutrophil population. Another possible hypothesis is that these neonatal mice start developing anti-rat antibodies to the depleting antibodies, but a 3-day period is too short for this to happen, and this effect would be seen in the circulating neutrophil compartment too, which we do not see.

The two major limitations of this protocol are outlined below. First, our study is restricted to neonatal mice. We chose P4 specifically as many neonatal studies on regulating the immune response are conducted at this age [14, 15]. Hence, we wanted to verify that adequate and complete neutrophil depletion can be achieved at this age. We urge investigators to test similar antibody combination strategies for experiments planned at other neonatal ages.

Further, our protocols did not precisely measure the other myeloid cell types. It has been well established that anti-Ly6G-1A8 clone antibody does not affect other myeloid cell lines and is specific for mature neutrophil populations. Our experiments were also carried out in C57Bl/6 mice, which do not express the gene for mouse IgG2a and hence cannot produce IgG2a subclass antibodies [26]. We encourage investigators using BALB/c and Swiss Webster mice, which produce IgG2a antibodies, to explore this protocol and verify the degree of neutrophil clearance before conducting experiments. In conclusion, a one-day administration of anti-Ly6G antibody followed after 4 hours by the anti-rat kappa immunoglobulin light chain antibody is extremely efficient in clearing neutrophils from the peripheral circulation, bone marrow, and spleen and should be verified with both intra- and extracellular Ly6G staining to determine the efficacy of neutrophil depletion.

## ACKNOWLEDGEMENTS

We acknowledge the use of the CWRU Cytometry Core for all flow cytometry experiments performed.

## Authors’ contributions

DM conceptualized experimental techniques, performed experiments, and wrote and reviewed the manuscript. SS and SC contributed to all experimental methods, flow cytometry interpretation, statistical analyses, and manuscript editing and reviewing. AT and YL contributed to experimental design and figure creation and reviewed and edited the manuscript. LN provided conceptual expertise and funding and reviewed and edited the manuscript.

